# Recovery of a distant large DNA virus from the genome sequencing of a fish leech

**DOI:** 10.64898/2026.11.15.766686

**Authors:** Alejandro Manzano-Marín, Anouk Willemsen

**Author notes:** Authors contributed equally to this work.

## Abstract

The leech gut is known to be a viral reservoir, where viruses can persist long-term and are protected from external influences. Therefore, leeches can serve as vectors for transmission of viral disease to susceptible hosts on which they prey. Nonetheless, viral diversity and virus-leech-host interactions remain largely unknown, with hitherto no research directed at identifying viruses that use leeches as their hosts. In the present study, we report the discovery of a novel large DNA virus (phylum *Nucleocytoviricota*) recovered from genomic sequencing data of the common fish leech *Piscicola geometra* (Linnaeus, 1761). Through genomic and phylogenetic analyses, we identify this lineage as highly divergent from currently known viruses, with ~95% of its predicted protein-coding genes showing no identifiable homology with current databases. Whether the leech is a host of this virus or a vector that transmits it to fish species remains to be investigated. This work highlights the value of off-target sequence retrieval for discovering novel viral lineages and sets the stage for future studies directed towards the prevalence and biology of leech-associated large DNA viruses.

## Introduction

Because of their haematophagous diet, and like other blood-feeding parasites, leeches are known carriers of disease agents, including protists, bacteria, and viruses (Burreson, 2006; CABI, 2019; Nehili *et al*., 1994). Although interactions between leeches and viruses have not been well characterised, leeches have been proposed as a potential viral reservoir for different viruses. Several studies show that mammalian viruses can persist long-term in the gut of *Hirudo medicinalis* leeches (Al-Khleif *et al*., 2011; Shope, 1957), suggesting that viruses are protected there from various deleterious chemical and physical influences to which they would be exposed in the open. These viruses include myxoma virus (dsDNA), equid alphaherpesvirus 1 (dsDNA), bovine parvovirus (ssDNA), classical swine fever virus (+ssRNA), feline calicivirus (+ssRNA), equine arteritis virus (+ssRNA) (Al-Khleif *et al*., 2011; Shope, 1957). Transmission of some of these viruses from the leech to susceptible hosts was also observed, highlighting a potential risk of using *H. medicinalis* leeches for medical purposes. Yet, leeches also offer a non-invasive resource for studying wildlife viral diversity and detecting novel, potentially zoonotic viruses before their emergence (Alfano *et al*., 2021; Kampmann *et al*., 2017).

Certain fish leeches have also been identified as virus vectors. The leech *Piscicola geometra* is a vector of spring viremia of carp virus (−ssRNA), leading to high mortality in carp pond cultures (Ahne, 1978, 1985). The leech *Piscicola salmositica* transmits infectious hematopoietic necrosis virus (−ssRNA) to salmonid fish, causing an acute rhabdoviral disease (Mulcahy *et al*., 1990). The leech *Myzobdella lugubris* may contribute to mortality in several freshwater fish species by transmitting viral hemorrhagic septicemia virus (−ssRNA) (Faisal and Schulz, 2009). Although the above-described fish leech-associated RNA viruses are classified within the same viral family, *Rhabdoviridae*, leeches are proposed to be a source of a large diversity of both DNA and RNA viruses (Alfano *et al*., 2021; Kampmann *et al*., 2017).

Large double-stranded DNA viruses within the phylum *Nucleocytoviricota* have attracted attention over the last decades, as this phylum includes viruses with exceptionally large virions and genomes (Abrahão *et al*., 2018; La Scola *et al*., 2003; Legendre *et al*., 2014; Philippe *et al*., 2013; Raoult *et al*., 2004), often referred to as giant viruses. Whereas many genes (~40-90%; Lücking *et al*., 2026) encoded in giant virus genomes remain functionally uncharacterised, these genomes also harbour components of the translation machinery and metabolic reprogramming pathways. A strain of the first discovered giant virus, *Acanthamoeba polyphaga* mimivirus, has been isolated from the internal organs and digestive tract of an *H. medicinalis* leech (Boughalmi *et al*., 2013). However, the role of leeches in hosting amoebae infecting giant viruses remains unclear.

Metagenomic sequencing of environmental samples from complex environments such as soil, marine water, freshwater, and wastewater treatment plants (Lücking *et al*., 2026; Minch and Moniruzzaman, 2025; Schulz *et al*., 2018, 2020; Zhang *et al*., 2015) has uncovered a large diversity of yet-uncultured *Nucleocytoviricota*. However, linking them to hosts or potential vectors remains challenging because of the complexity of the host community in the sample and technical limitations in sequencing of the eukaryotic fraction. Recent large-scale sequencing projects have generated high-quality eukaryotic genomes, including the *European Reference Genome Atlas* (Mazzoni *et al*., 2023), *Vertebrate Genomes Project* (Rhie *et al*., 2021), *Aquatic Symbiosis Genomics Project* (McKenna *et al*., 2024), *Darwin Tree of Life* (The Darwin Tree of Life Project Consortium *et al*., 2022), among others (Blaxter *et al*., 2025). The sequencing data from these projects represent an untapped resource for uncovering co-inhabiting entities, such as symbiotic microbes and viruses.

In this work, we report the serendipitous recovery of an abundant large double-stranded DNA virus, hereafter “NCV PIGEOM”, from the genome sequencing data from the fish leech *P. geometra* (Doe *et al*., 2023). Genomic analysis revealed that almost all predicted protein-coding genes lack homology to sequences in currently available public databases, mirroring the divergence observed with other *Nucleocytoviricota* in phylogenetic reconstructions using different hallmark genes. This study highlights the utility and potential for discoveries from careful analysis and curation of off-target sequenced entities. Further, our work highlights leeches as potential reservoirs of large DNA viruses that can be transmitted to different fish species, expanding the range of known invertebrate interactions.

## Results & Discussion

### Genome of a leech-associated large DNA virus

We retrieved PacBio HiFi reads from the chromosome-level reference genome of the fish leech *P. geometra* (Linnaeus, 1761) (Annelida: Clitellata: Euhirudinea: Oceanobdelliformes) from the SRA archive, generated as part of the *Darwin Tree of Life project* (Doe *et al*., 2023). We performed *de novo* assembly with Flye and recovered a circular molecule of 204,626 Mega base-pairs, with an average coverage of 89x (compared to 38x for the leech scaffolds) and an average GC content of 21.93%. Visual inspection of the assembly graphs revealed no connections to any other sequence in the assembly. A prediction of rRNA gene sequences using rnammer returned no hits, suggesting a plasmidic or non-bacterial origin. An initial draft annotation of the molecule using Prokka identified 115 putative protein-coding genes, none with a functional annotation. A subsequent search of predicted coding sequences using InterProScan returned a DNA-directed DNA polymerase, family B (PolB). BLASTP analysis of the PolB sequence using the NCBI web server showed best hits against several viruses within the phylum *Nucleocytoviricota*, including those within the *Iridoviridae, Ascoviridae*, and *Pithoviridae* families (**Table 1**). Because of its suspected viral origin, we ran the MMSeqs2-2 pipeline (Camargo, 2026; Lood *et al*., 2026) to infer its affiliation. The results further supported its assignment to *Nucleocytoviricota* (phylum score=0.7013). Our gene prediction and annotation approaches failed to detect the typical double jelly-roll major capsid protein (MCP), a viral hallmark gene for most *Nucleocytoviricota* (Krupovič and Bamford, 2008; Krupovic and Koonin, 2026). Thus, this virus likely encodes either a highly divergent homologous MCP or has evolutionarily lost the ancestral MCP, which was replaced by unrelated proteins, as in pithoviruses and pandoraviruses, resulting in odd capsid shapes (Krupovic and Koonin, 2026). Future studies that include virus isolation combined with electron microscopy will allow us to assess the virion morphology of “NCV PIGEOM”.

**Table 1.** Top 10* BLASTP hits of PolB *vs*. the ClusteredNR database. * Three bacterial hits were removed due to suspected contamination or wrong assignment, as they originated from highly fragmented Metagenome-Assembled Genomes (MAGs) with no downstream curation. ^M^ MAG. ^I^ Isolate. ^R^ Complete CDS from RNA-Seq experiment.

| Family | Accession | Query coverage (%) | E-value | identity (%) | Description |
|---|---|---|---|---|---|
| *Iridoviridae* | DBA45601.1 | 66 | 9e-35 | 24.47 | Iridovirus macaca37137<sup>M</sup> |
|  | YP_009163831.1 | 56 | 7e-33 | 24.03 | *Megalocytivirus lates1* (=Scale drop disease virus)<sup>I</sup> |
|  | XBE43076.1 | 67 | 6e-29 | 22.83 | Chrysiptera rollandi iridovirus<sup>M</sup> |
|  | QXT57784.1 | 60 | 1e-28 | 24.04 | Rhinella marina erythrocytic-like virus<sup>R</sup> |
| *Pithoviridae* | QBK90645.1 | 61 | 6e-31 | 25.18 | *Pithovirus* LCPAC104<sup>M</sup> |
|  | GMQ13763.1 | 69 | 7e-31 | 23.12 | *Cedratvirus* Ce7-1<sup>I</sup> |
|  | YP_009000951.1 | 77 | 1e-29 | 22.61 | *Alphapithovirus siberiense* (=*Pithovirus sibericum*)<sup>I</sup> |
|  | YP_013022960.1 | 61 | 7e-29 | 23.81 | *Alphacedratvirus rossiense* (=’Cedratvirus kamchatka’)<sup>I</sup> |
|  | WIL04983.1 | 61 | 9e-29 | 23.81 | *Alphacedratvirus rossiense* (=’Cedratvirus duvanny’)<sup>I</sup> |
| *Ascoviridae* | XNL98089.1 | 54 | 9e-30 | 25.83 | Androctonus mauritanicus toursvirus<sup>M</sup> |

### Phylogenetic analysis places the virus within *Pimascovirales*

Consistent with the draft annotation, we initially identified only one *Nucleocytoviricota* marker gene (PolB) in the “NCV PIGEOM” virus genome using HMM model searches. However, using Foldseek, we identified four additional marker genes: SFII, RNAPS, RNAPL, and TopoII. We aligned these sequences with those extracted from 271 reference genomes in the *Giant Virus Database* (GVDB), resulting in 232 taxa for SFII, 209 for RNAPS, 201 for RNAPL, 267 for PolB, and 201 for TopoII. We filtered the alignments and performed phylogenetic reconstruction on the individual and concatenated protein alignments. Model selection indicated LG+F+I+G4 as the best model for SFII, RNAPS, RNAPL, and PolB, and Q.PFAM+F+I+G4 for TopoII. Individual phylogenetic analyses for 3/5 marker genes placed “NCV PIGEOM” within the *Pimascovirales* order (SFII, PolB, TopoII), whereas for the other 2/5 marker genes “NCV PIGEOM” was placed either basal to the *Pimascovirales* (RNAPS) or placed between the outgroup *Chitovirales* and all other currently recognised viral orders within *Nucleocytoviricota* (RNAPL) (supplementary **Figures S1-S5**). Phylogenetic analysis of the concatenated alignments (272 taxa) supported placement within the *Pimascovirales*, basal to *Ocovirineae*, the clade containing pithoviruses, cedratviruses and orpheovirus, with an ML bootstrap support of 97.9 (**Figure 1**). This phylogenetic positioning supports the failure to detect an MCP gene, as *Ocovirineae* have evolutionarily replaced this gene (Claverie *et al*., 2025). Performing the same analysis (including realignment and alignment filtering) but keeping only taxa with all five core genes present in the dataset (135 taxa), we obtained the same positioning (supplementary **Figure S6**). We also recovered the placement using only the *Pimascovirales* (**Figure 2**). The long branch leading to “NCV PIGEOM” indicates that this taxon is very distant from other taxa within the *Pimascovirales*, and finding more genomes similar to this new viral genome might support classifying “NCV PIGEOM” within a novel viral group.

**Figure 1.**
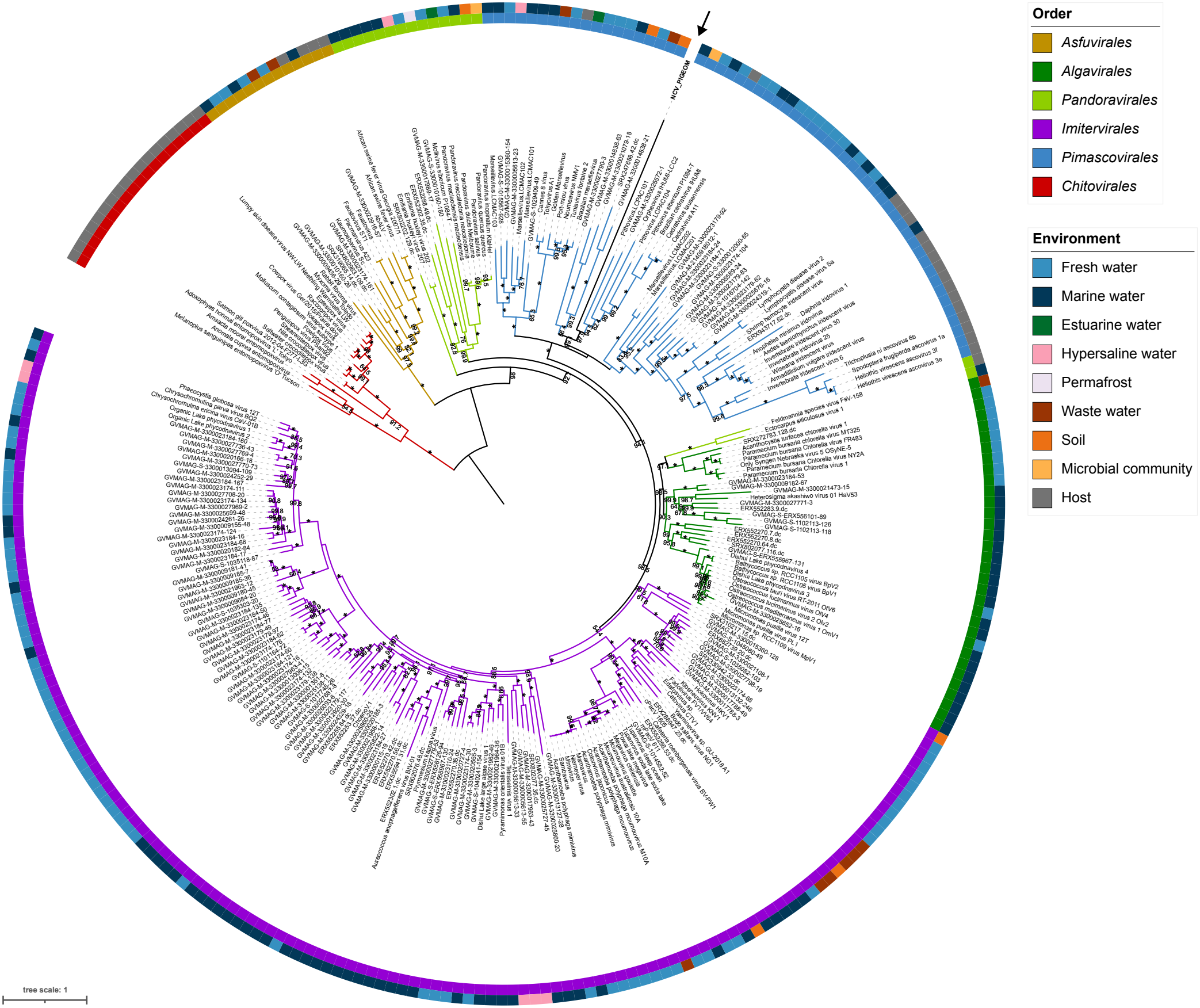
Maximum likelihood phylogenetic tree of 272 *Nucleocytoviricota*. The tree was constructed at the amino acid level from concatenated alignments of the *SFII, RNAPS, RNAPL, PolB*, and *TopoII* core genes. Branch values correspond to bootstrap support values between 50 and 100. Bootstrap support of 100 is represented with an asterisk. *Chitovirales* was used as the outgroup. Branch colour coding and colours in the first ring around the tree correspond to the viral orders. Colours in the second ring correspond to the environment in which the virus was found. The arrow indicates the location of “NCV PIGEOM”.

**Figure 2.**
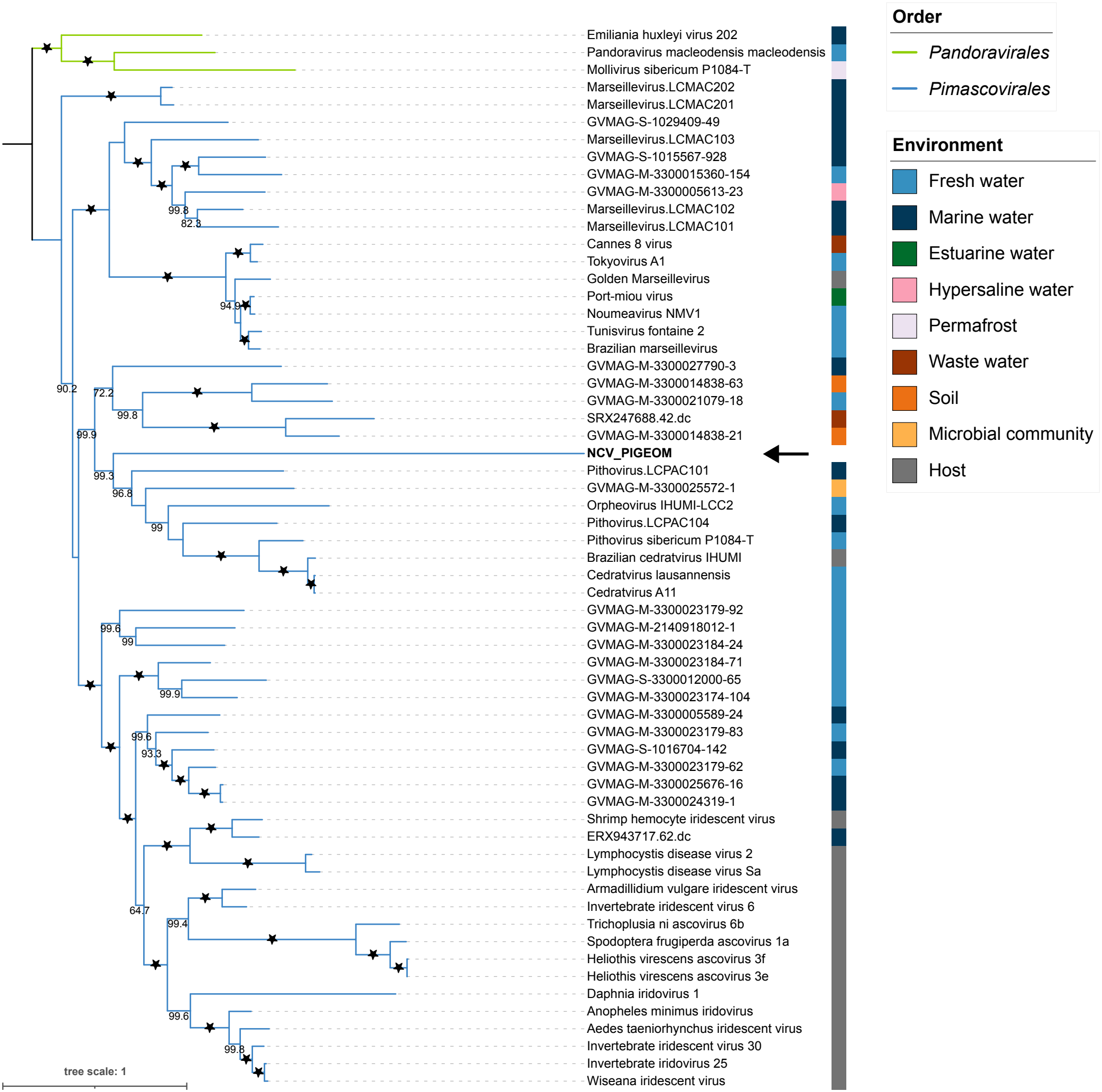
Maximum likelihood phylogenetic tree for representatives of the *Pimascovirales* order. The tree was constructed at the amino acid level from concatenated alignments of the *SFII, RNAPS, RNAPL, PolB*, and *TopoII* core genes from the “NCV PIGEOM”, 60 pimascovirid and 3 pandoravirid genomes. Branch values correspond to bootstrap support values between 50 and 100. Bootstrap support of 100 is represented with a black star. Branch colour coding corresponds to the viral orders. Members of *Pandoravirales* were used as the outgroup. Colours on the right correspond to the environment in which the virus was found. The arrow indicates the location of “NCV PIGEOM”.

### *P. geometra*, a vector or a host of *Nucleocytoviricota*?

The apparent high coverage of “NCV PIGEOM” relative to the host genome suggests this virus was abundant within the sequenced leech. However, in the original sequencing study, no tissue dissection was performed to separate the digestive tract from leftover tissue, leaving the question open as to whether the leech can be formally proposed as a vector or host for this newly identified large DNA virus. As described in the introduction, different fish leeches have been proposed to transmit different viruses, often associated with disease and high mortality rates in different fish species (Ahne, 1978, 1985; Faisal and Schulz, 2009; Mulcahy *et al*., 1990). However, none of these cases demonstrated that the viruses infected leech body tissues, nor did they show viruses replicating in the gut, supporting the role of leeches as pure vectors by providing a protective environment for transmission.

Our BLASTP (**Table 1**) and phylogenetic analyses (**Figures 1**-**2**, and supplementary **Figures S1-S6**) suggest affiliation with viruses from the *Ocovirineae* suborder, and *Ascoviridae* and *Iridoviridae* families. Currently reported *Ocovirineae* isolates infect amoebae (*Acanthamoeba* spp. and *Vermamoeba vermiformis*), but other strains have also been recovered from environmental metagenomic sequencing datasets, with no identified hosts (Claverie *et al*., 2025). Members of the *Ascoviridae* family mainly infect lepidopteran larvae and pupae, and are mechanically transmitted by parasitoid wasps (Bigot *et al*., 1997; Federici *et al*., 2009; Glynn Tillman *et al*., 2004). *Iridoviridae* is currently classified into two subfamilies, *Alphairidovirinae* and *Betairidovirinae*. The *Alphairidovirinae* contains viruses that primarily infect ectothermic vertebrates such as bony fish, amphibians and reptiles, whereas *Betairidovirinae* mainly infect invertebrates such as insects and crustaceans. Comparative genomics combined with phylogenetics shows several recent host shifts among iridoviruses, with those infecting fish being ancestral to those infecting amphibians (Jancovich *et al*., 2010). The long branch leading to “NCV PIGEOM” and the observed differences in phylogenetic positioning depending on the core gene used support these observations. However, the lack of homology with other fish iridoviruses and lack of information on viruses infecting the fish in the same environment where the leech was sampled do not allow us to assign “NCV PIGEOM” to a fish host.

Due to the information gap on the biology and natural host of this virus, we cannot completely exclude the possibility that “NCV PIGEOM” infects the leech *P. geometra*. The high abundance and long-term stability of African swine fever virus in the leech *H. medicinalis* led researchers to investigate this possibility. Indeed, viral particles adhered to the leech skin, and early viral genes were expressed in leech body cells; however, differences in early- and late-gene expression and the lack of infectious viral particle production indicated abortive infections (Avagyan *et al*., 2023). Whether this is also the case for “NCV PIGEOM” remains to be investigated. Future studies that systematically detect and isolate large DNA viruses in different leech tissues (including ingested blood) across populations using molecular and microscopy techniques will shed light on this aspect. Subsequent studies on the biology of these isolated viruses will clarify the relationship between this novel virus and will reveal its potential vectors and hosts.

## Materials and Methods

### Genome assembly and annotation

We downloaded PacBio Hi-Fi sequencing reads from the leech *P. geometra* from the NCBI SRA database (accession SRS4575368). Reads were then assembled de novo using Flye v2.9.2 (Kolmogorov *et al*., 2019) using the ‘–pacbio-hifi’ mode and two polishing iterations. We predicted rRNA gene sequences using rnammer v1.2 (Lagesen *et al*., 2007). We reoriented the molecule by predicting the origin of replication using OriginX (Worning *et al*., 2006). Gene prediction and draft genome annotation were performed using Prokka v1.14.6 (Seemann, 2014). Further functional and domain predictions were performed using InterProScan v5.78-109.0 (Jones *et al*., 2014). We performed RNA gene and motif prediction using Infernal v1.1.5 (Nawrocki and Eddy, 2013) against the Rfam database v15.1 (Ontiveros-Palacios *et al*., 2025). After performing BLAST-based searches, we assigned sequences to virus taxa using MMseqs2 (Mirdita *et al*., 2021) with a database of virus proteins annotated with taxonomic data, following the MMSeqs2-2 pipeline (Camargo, 2026), which has been shown to perform well for characterising sequences from unexplored viromes (Lood *et al*., 2026). Scores for a taxon in this pipeline are calculated as the sum of the weights of the ORFs that are assigned to it, divided by the sum of the weights of all ORFs.

### Phylogenetic reconstruction

First, we downloaded 271 *Nucleocytoviricota* reference genomes from the Giant Virus Database (GVDB: https://faylward.github.io/GVDB/, last accessed November 13, 2026). We used ncldv markersearch (Moniruzzaman *et al*., 2020) and HMMER v3.3.2 (http://hmmer.org/, last accessed November 13, 2026) for the identification of nine phylogenetic marker genes (MCP: major capsid protein, SFII: DEAD/SNF2-like helicase, RNAPS: DNA-directed RNA polymerase beta subunit, RNAPL: DNA-directed RNA polymerase alpha subunit, PolB: DNA polymerase family B, TFIIB: Transcription initiation factor IIB, TopoII: DNA topoisomerase II, A32: A32-like Packaging ATPase, VLTF3: Poxvirus Late Transcription Factor VLTF3). Because this method identified only one marker gene (PolB) in the “NCV PIGEOM” genome, we used Foldseek v10-941cd33-gpu (Van Kempen *et al*., 2024) to identify additional marker genes. In brief, we created reference databases for the nine marker proteins using the viral genomes extracted from the GVDB and a target database for all predicted proteins in the “NCV PIGEOM” genome, then used the easy-search module to query the reference protein sequences against the target database. The amino acid sequences of five identified (SFII, RNAPS, RNAPL, PolB and TopoII) marker genes were aligned with MAFFT v7.526 (Katoh and Standley, 2013) using the G-INS-i method and filtered using Gblocks (Castresana, 2000) with the parameters ‘-t=p -b1=50%+1 -b2=50%+1 -b3=8 -b4=2 -b5=a -b0=2’. The alignments were concatenated with catfasta2phyml v1.2.1 (https://github.com/nylander/catfasta2phyml, last accessed November 13, 2026). We performed maximum likelihood phylogenetic analyses on the individual and concatenated protein alignments using IQ-TREE v3.1.0 (Wong *et al*., 2026), including ModelFinder (Kalyaanamoorthy *et al*., 2017) for model selection, the ultrafast bootstrap approximation (Hoang *et al*., 2018; Minh *et al*., 2013), and the SH-like approximate likelihood ratio test (Guindon *et al*., 2010). For the concatenated alignments, we used partitions to indicate the appropriate model for each protein sequence.

## Supplementary Material & Data Availability

Supplementary **Figures S1-S6**, the “NCV PIGEOM” genome, annotation files, Foldseek results, alignments, and phylogenetic trees are available at the Zenodo repository 10.5281/zenodo.21997283. The newly sequenced and annotated genome is in the process of being accessioned at the European Nucleotide Archive (ENA).

## Acknowledgements

The computational results of this work have been achieved using the *Life Science Compute Cluster* (LiSC) of the University of Vienna. This project has received funding from the European Union (ERC, CHIMERA, 101039843 to AW). Views and opinions expressed are, however, those of the author(s) only and do not necessarily reflect those of the European Union or the *European Research Council Executive Agency*. Neither the European Union nor the granting authority can be held responsible for them. This research was funded in whole or in part by the *Austrian Science Fund* (FWF) 10.55776/TAI1450724 to AMM and 10.55776/COE7 to AW.

## References

Abrahão J, Silva L, Silva L. S, Khalil J. Y. B, Rodrigues R, Arantes T, Assis F, Boratto P, Andrade M, Kroon E. G, Ribeiro B, Bergier I, Seligmann H, Ghigo E, et al. 2018. Tailed giant Tupanvirus possesses the most complete translational apparatus of the known virosphere. Nature Communications, 9(1): 749. URL 10.1038/s41467-018-03168-1.

Ahne W. 1978. Uptake and multiplication of spring viraemia of carp virus in carp, Cyprinus carpio L. Journal of Fish Diseases, 1(3): 265–268. URL 10.1111/j.1365-2761.1978.tb00029.x.

Ahne W. 1985. Argulus foliaceus l. and Piscicola geometra l. as mechanical vectors of spring viraemia of carp virus (svcv). Journal of Fish Diseases, 8(2): 241–242. URL 10.1111/j.1365-2761.1985.tb01220.x.

Al-Khleif A, Roth M, Menge C, Heuser J, Baljer G, and Herbst W. 2011. Tenacity of mammalian viruses in the gut of leeches fed with porcine blood. Journal of Medical Microbiology, 60(6): 787–792. URL 10.1099/jmm.0.027250-0.

Alfano N, Dayaram A, Axtner J, Tsangaras K, Kampmann M.-L, Mohamed A, Wong S. T, Gilbert M. T. P, Wilting A, and Greenwood A. D. 2021. Non-invasive surveys of mammalian viruses using environmental dna. Methods in Ecology and Evolution, 12(10): 1941–1952. URL 10.1111/2041-210X.13661.

Avagyan H. R, Hakobyan S. A, Avetisyan A. S, Bayramyan N. V, Hakobyan L. H, Poghosyan A. A, Abroyan L. O, Baghdasaryan B. V, Tsakanova G. V, Sahakyan L. V, Yeremyan A. S, and Karalyan Z. A. 2023. The pattern of stability of african swine fever virus in leeches. Veterinary Microbiology, 284: 109835. URL 10.1016/j.vetmic.2023.109835.

Bigot Y, Periquet G, Sizaret P. Y, Doury G, Delbost F, Hamelin M. H, and Rabouille A. 1997. Biological and molecular features of the relationships between diadromus pulchellus ascovirus, a parasitoid hymenopteran wasp (Diadromus pulchellus) and its lepidopteran host, Acrolepiopsis Assectella. Journal of General Virology, 78(5): 1149–1163. URL 10.1099/0022-1317-78-5-1149.

Blaxter M, Lewin H. A, DiPalma F, Challis R, Da Silva M, Durbin R, Formenti G, Franz N, Guigo R, Harrison P. W, Hiller M, Hoff K. J, Howe K, Jarvis E. D, et al. 2025. The Earth BioGenome Project phase II: Illuminating the eukaryotic tree of life. Frontiers in Science, 3: 1514835. URL 10.3389/fsci.2025.1514835.

Boughalmi M, Pagnier I, Aherfi S, Colson P, Raoult D, and La Scola B. 2013. First isolation of a giant virus from wild Hirudo medicinalis leech: Mimiviridae isolation in Hirudo medicinalis. Viruses, 5(12): 2920–2930. URL 10.3390/v5122920.

Burreson E. M. 2006. Phylum annelida: Hirudinea as vectors and disease agents. In P. T. K. Woo, editor, Fish diseases and disorders. Volume 1: Protozoan and metazoan infections, pages 566–591. CABI, UK, 2 edition. URL 10.1079/9780851990156.0566.

CABI. 2019. Hirudinea as vectors and disease agents in fish. CABI Compendium, Article no. 79600. URL 10.1079/cabicompendium.79600.

Camargo A. 2026. Apcamargo/ictv-mmseqs2-protein-database [release vmr msl39.v4 20241106 v2]. URL https://github.com/apcamargo/ictv-mmseqs2-protein-database.

Castresana J. 2000. Selection of conserved blocks from multiple alignments for their use in phylogenetic analysis. Molecular Biology and Evolution, 17(4): 540–52. URL 10.1093/oxfordjournals.molbev.a026334.

Claverie J.-M, Legendre M, Rigou S, and Abergel C. 2025. Refining the taxonomy of Pithovirus-related giant dna viruses within the order Pimascovirales. Archives of Virology, 170(5): 111. URL 10.1007/s00705-025-06297-3.

Doe J, Lab N. H. M. G. A, Darwin Tree of Life Barcoding collective, Wellcome Sanger Institute Tree of Life programme, Wellcome Sanger Institute Scientific Operations: DNA Pipelines collective, Tree of Life Core Informatics collective, and Darwin Tree of Life Consortium. 2023. The genome sequence of the fish leech, Piscicola geometra (linnaeus, 1761). Wellcome Open Research, 8: 229. URL 10.12688/wellcomeopenres.19488.1.

Faisal M and Schulz C. A. 2009. Detection of viral hemorrhagic septicemia virus (vhsv) from the leech Myzobdella lugubris leidy, 1851. Parasites & Vectors, 2(1): 45. URL 10.1186/1756-3305-2-45.

Federici B. A, Bideshi D. K, Tan Y, Spears T, and Bigot Y. 2009. Ascoviruses: Superb manipulators of apoptosis for viral replication and transmission. In J. L. Van Etten, R. W. Compans, M. D. Cooper, T. Honjo, H. Koprowski, F. Melchers, M. B. A. Oldstone, S. Olsnes, and P. K. Vogt, editors, Lesser known large dsDNA viruses, volume 328 of Current topics in microbiology and immunology, pages 171–196. Springer Berlin Heidelberg, Berlin, Heidelberg. URL 10.1007/978-3-540-68618-7_5.

Glynn Tillman P, Styer E. L, and Hamm J. J. 2004. Transmission of Ascovirus from Heliothis virescens (lepidoptera: Noctuidae) by three parasitoids and effects of virus on survival of parasitoid Cardiochiles nigriceps (hymenoptera: Braconidae). Environmental Entomology, 33(3): 633–643. URL 10.1603/0046-225X-33.3.633.

Guindon S, Dufayard J. F, Lefort V, Anisimova M, Hordijk W, and Gascuel O. 2010. New algorithms and methods to estimate maximum-likelihood phylogenies: assessing the performance of phyml 3.0. Systematic Biology, 59(3): 307–321. URL 10.1093/sysbio/syq010.

Hoang D. T, Chernomor O, von Haeseler A, Minh B. Q, and Vinh L. S. 2018. Ufboot2: Improving the ultrafast bootstrap approximation. Molecular Biology and Evolution, 35(2): 518–522. URL 10.1093/molbev/msx281.

Jancovich J. K, Bremont M, Touchman J. W, and Jacobs B. L. 2010. Evidence for multiple recent host species shifts among the ranaviruses (family Iridoviridae). Journal of Virology, 84(6): 2636–2647. URL 10.1128/JVI.01991-09.

Jones P, Binns D, Chang H.-Y, Fraser M, Li W, McAnulla C, McWilliam H, Maslen J, Mitchell A, Nuka G, Pesseat S, Quinn A. F, Sangrador-Vegas A, Scheremetjew M, et al. 2014. Interproscan 5: Genome-scale protein function classification. Bioinformatics, 30(9): 1236–1240. URL 10.1093/bioinformatics/btu031.

Kalyaanamoorthy S, Minh B. Q, Wong T. K. F, von Haeseler A, and Jermiin L. S. 2017. Modelfinder: Fast model selection for accurate phylogenetic estimates. Nature Methods, 14(6): 587–589. URL 10.1038/nmeth.4285.

Kampmann M.-L, Schnell I. B, Jensen R. H, Axtner J, Sander A. F, Hansen A. J, Bertelsen M. F, Greenwood A. D, Gilbert M. T. P, and Wilting A. 2017. Leeches as a source of mammalian viral dna and rna—a study in medicinal leeches. European Journal of Wildlife Research, 63(2): 36. URL 10.1007/s10344-017-1093-6.

Katoh K and Standley D. M. 2013. Mafft multiple sequence alignment software version 7: Improvements in performance and usability. Molecular Biology and Evolution, 30(4): 772–780. URL 10.1093/molbev/mst010.

Kolmogorov M, Yuan J, Lin Y, and Pevzner P. A. 2019. Assembly of long, error-prone reads using repeat graphs. Nature Biotechnology, 37(5): 540–546. URL 10.1038/s41587-019-0072-8.

Krupovič M and Bamford D. H. 2008. Virus evolution: How far does the double β-darrel viral lineage extend? Nature Reviews Microbiology, 6(12): 941–948. URL 10.1038/nrmicro2033.

Krupovic M and Koonin E. V. 2026. Organization and evolution of the virosphere and the replicator space. Comptes Rendus. Biologies, 349(G1): 35–75. URL 10.5802/crbiol.193.

La Scola B, Audic S, Robert C, Jungang L, de Lamballerie X, Drancourt M, Birtles R, Claverie J.-M, and Raoult D. 2003. A giant virus in amoebae. Science, 299(5615): 2033. URL 10.1126/science.1081867.

Lagesen K, Hallin P, Rodland E. A, Staerfeldt H. H, Rognes T, and Ussery D. W. 2007. Rnammer: Consistent and rapid annotation of ribosomal rna genes. Nucleic Acids Res, 35(9): 3100–3108. URL 10.1093/nar/gkm160.

Legendre M, Bartoli J, Shmakova L, Jeudy S, Labadie K, Adrait A, Lescot M, Poirot O, Bertaux L, Bruley C, Coute Y, Rivkina E, Abergel C, Claverie J.-M. J.-M, et al. 2014. Thirty-thousand-year-old distant relative of giant icosahedral dna viruses with a Pandoravirus morphology. Proceedings of the National Academy of Sciences of the United States of America, 111(11): 4274–4279. URL 10.1073/pnas.1320670111.

Lood C, Doijad S. P, Adriaenssens E. M, Bao Y, Barylski J, Bolduc B, Bouras G, Brister J. R, Brown C. T, Camargo A. P, Coninck L. D, Deorowicz S, Edgar R, Edwards R. A, et al. 2026. First community challenge for automated virus taxonomy. bioRxiv, page 2026.07.04.736517. URL 10.64898/2026.07.04.736517.

Lücking D, Manzano-Marín A, and Willemsen A. 2026. Multilayered network analysis reveals the centrality of newly discovered Nucleocytoviricota in wastewater treatment plant communities. The ISME Journal, page wrag267. URL 10.1093/ismejo/wrag267.

Mazzoni C. J, Ciofi C, and Waterhouse R. M. 2023. Biodiversity: An atlas of european reference genomes. Nature, 619(7969): 252–252. URL 10.1038/d41586-023-02229-w.

McKenna V, Archibald J, Beinart R, Dawson M, Hentschel U, Keeling P, Lopez J, Martín-Durán J, Petersen J, Sigwart J, Simakov O, Sutherland K, Sweet M, Talbot N, et al. 2024. The Aquatic Symbiosis Genomics Project: Probing the evolution of symbiosis across the Tree of Life. Wellcome Open Research, 6(254). URL 10.12688/wellcomeopenres.17222.2.

Minch B and Moniruzzaman M. 2025. Expansion of the genomic and functional diversity of global ocean giant viruses. npj Viruses, 3(1): 32. URL 10.1038/s44298-025-00122-z.

Minh B. Q, Nguyen M. A. T, and Von Haeseler A. 2013. Ultrafast Approximation for Phylogenetic Bootstrap. Molecular Biology and Evolution, 30(5): 1188–1195. URL 10.1093/molbev/mst024.

Mirdita M, Steinegger M, Breitwieser F, Söding J, and Levy Karin E. 2021. Fast and sensitive taxonomic assignment to metagenomic contigs. Bioinformatics, 37(18): 3029–3031. URL 10.1093/bioinformatics/btab184.

Moniruzzaman M, Martinez-Gutierrez C. A, Weinheimer A. R, and Aylward F. O. 2020. Dynamic genome evolution and complex virocell metabolism of globally-distributed giant viruses. Nature Communications, 11(1): 1710. URL 10.1038/s41467-020-15507-2.

Mulcahy D. M, Klaybor D, and Batts W. N. 1990. Isolation of infectious hematopoietic necrosis virus from a leech (Piscicola salmositica) and a copepod (Salmincola sp.), ectoparasites of sockeye salmon Oncorhynchus nerka. Diseases of Aquatic Organisms, 8: 29–34. URL 10.3354/dao008029.

Nawrocki E. P and Eddy S. R. 2013. Infernal 1.1: 100-fold faster RNA homology searches. Bioinformatics, 29(22): 2933–2935. URL 10.1093/bioinformatics/btt509.

Nehili M, Ilk C, Mehlhorn H, Ruhnau K, Dick W, and Njayou M. 1994. Experiments on the possible role of leeches as vectors of animal and human pathogens: A light and electron microscopy study. Parasitology Research, 80(4): 277–290. URL 10.1007/BF02351867.

Ontiveros-Palacios N, Cooke E, Nawrocki E. P, Triebel S, Marz M, Rivas E, Griffiths-Jones S, Petrov A. I, Bateman A, and Sweeney B. 2025. Rfam 15: RNA families database in 2025. Nucleic Acids Research, 53(D1): D258–D267. URL 10.1093/nar/gkae1023.

Philippe N, Legendre M, Doutre G, Coute Y, Poirot O, Lescot M, Arslan D, Seltzer V, Bertaux L, Bruley C, Garin J, Claverie J.-M, and Abergel C. 2013. Pandoraviruses: Amoeba viruses with genomes up to 2.5 Mb reaching that of parasitic eukaryotes. Science, 341(6143): 281–286. URL 10.1126/science.1239181.

Raoult D, Audic S, Robert C, Abergel C, Renesto P, Ogata H, La Scola B, Suzan M, and Claverie J.-M. 2004. The 1.2-megabase genome sequence of Mimivirus. Science, 306(5700): 1344–1350. URL 10.1126/science.1101485.

Rhie A, McCarthy S. A, Fedrigo O, Damas J, Formenti G, Koren S, Uliano-Silva M, Chow W, Fungtammasan A, Kim J, Lee C, Ko B. J, Chaisson M, Gedman G. L, et al. 2021. Towards complete and error-free genome assemblies of all vertebrate species. Nature, 592(7856): 737–746. URL 10.1038/s41586-021-03451-0.

Schulz F, Alteio L, Goudeau D, Ryan E. M, Yu F. B, Malmstrom R. R, Blanchard J, and Woyke T. 2018. Hidden diversity of soil giant viruses. Nature Communications, 9(1): 4881. URL 10.1038/s41467-018-07335-2.

Schulz F, Roux S, Paez-Espino D, Jungbluth S, Walsh D, Denef V. J, McMahon K. D, Konstantinidis K. T, Eloe-Fadrosh E. A, Kyrpides N, and Woyke T. 2020. Giant virus diversity and host interactions through global metagenomics. Nature. URL 10.1038/s41586-020-1957-x.

Seemann T. 2014. Prokka: Rapid prokaryotic genome annotation. Bioinformatics, 30(14): 2068–2069. URL 10.1093/bioinformatics/btu153.

Shope R. E. 1957. The leech as a potential virus reservoir. The Journal of Experimental Medicine, 105(4): 373–382. URL 10.1084/jem.105.4.373.

The Darwin Tree of Life Project Consortium, Blaxter M, Mieszkowska N, Di Palma F, Holland P, Durbin R, Richards T, Berriman M, Kersey P, Hollingsworth P, Wilson W, Twyford A, Gaya E, Lawniczak M, et al. 2022. Sequence locally, think globally: The Darwin Tree of Life Project. Proceedings of the National Academy of Sciences, 119(4): e2115642118. URL 10.1073/pnas.2115642118.

Van Kempen M, Kim S. S, Tumescheit C, Mirdita M, Lee J, Gilchrist C. L. M, Söding J, and Steinegger M. 2024. Fast and accurate protein structure search with Foldseek. Nature Biotechnology, 42(2): 243–246. URL 10.1038/s41587-023-01773-0.

Wong T. K. F, Ly-Trong N, Ren H, Demotte P, Baños H, Roger A. J, Susko E, Bielow C, De Maio N, Goldman N, Hahn M. W, Dos Reis M, Vinh L. S, Huttley G, et al. 2026. IQ-TREE 3: Phylogenomic inference software using complex evolutionary models. Molecular Biology and Evolution, 43(5): msag117. URL 10.1093/molbev/msag117.

Worning P, Jensen L. J, Hallin P. F, Stærfeldt H.-H, and Ussery D. W. 2006. Origin of replication in circular prokaryotic chromosomes. Environmental Microbiology, 8(2): 353–361. URL 10.1111/j.1462-2920.2005.00917.x.

Zhang W, Zhou J, Liu T, Yu Y, Pan Y, Yan S, and Wang Y. 2015. Four novel algal virus genomes discovered from Yellowstone Lake metagenomes. Scientific Reports, 5(1): 15131. URL 10.1038/srep15131.

