## Supplementary figures S1-S6 for "Recovery of a distant large DNA virus from the genome sequencing of a fish leech"

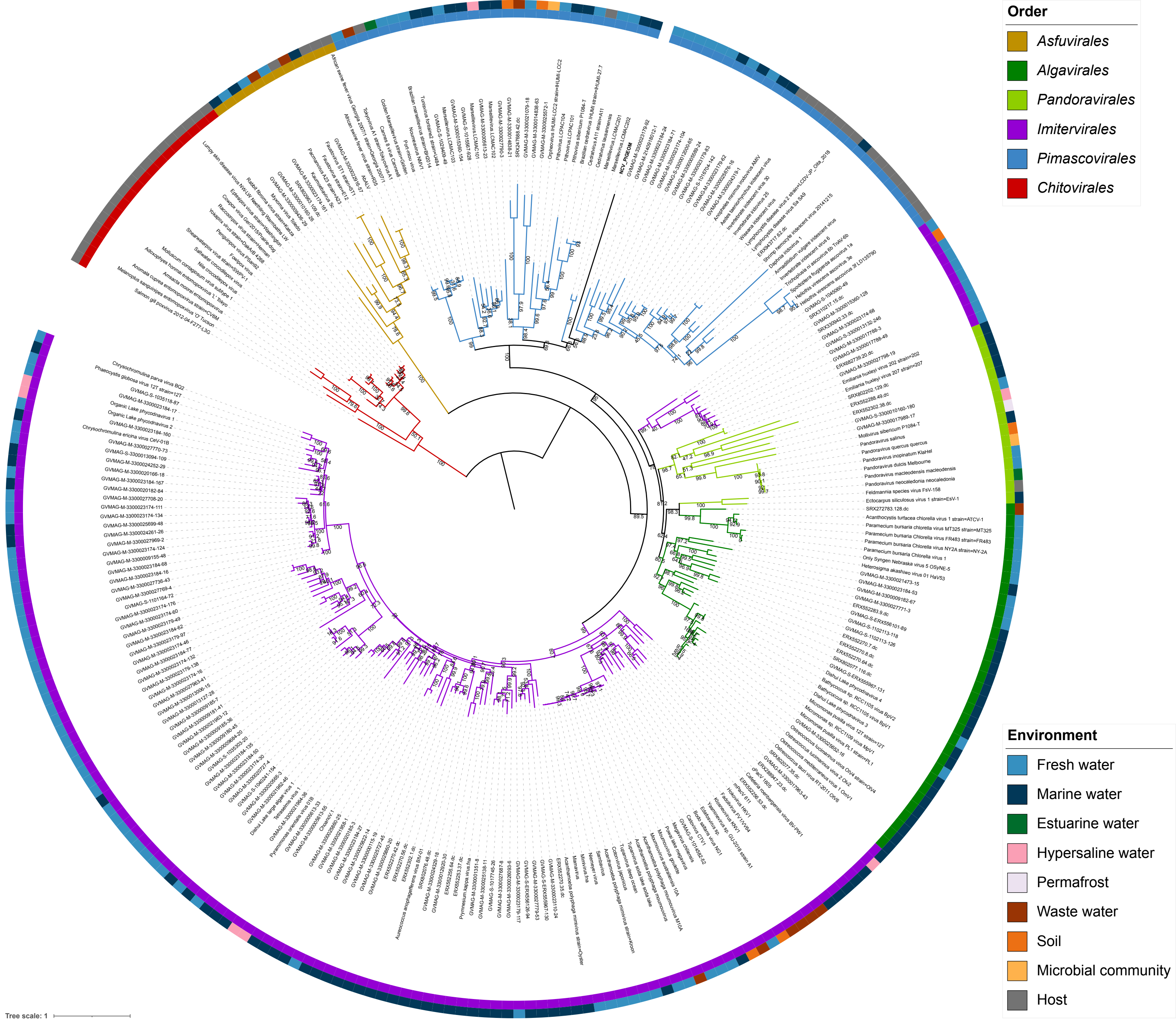


**Figure S1**. Maximum likelihood phylogenetic tree for *PolB* with a dataset containing 267 *Nucleocytoviricota*. The tree was constructed at the amino acid level. Branch values correspond to bootstrap support values. Branch colour coding and colours in the first ring around the tree correspond to the viral orders. *Chitovirales* was used as the outgroup. The colours in the second ring correspond to the environment in which the virus was found.

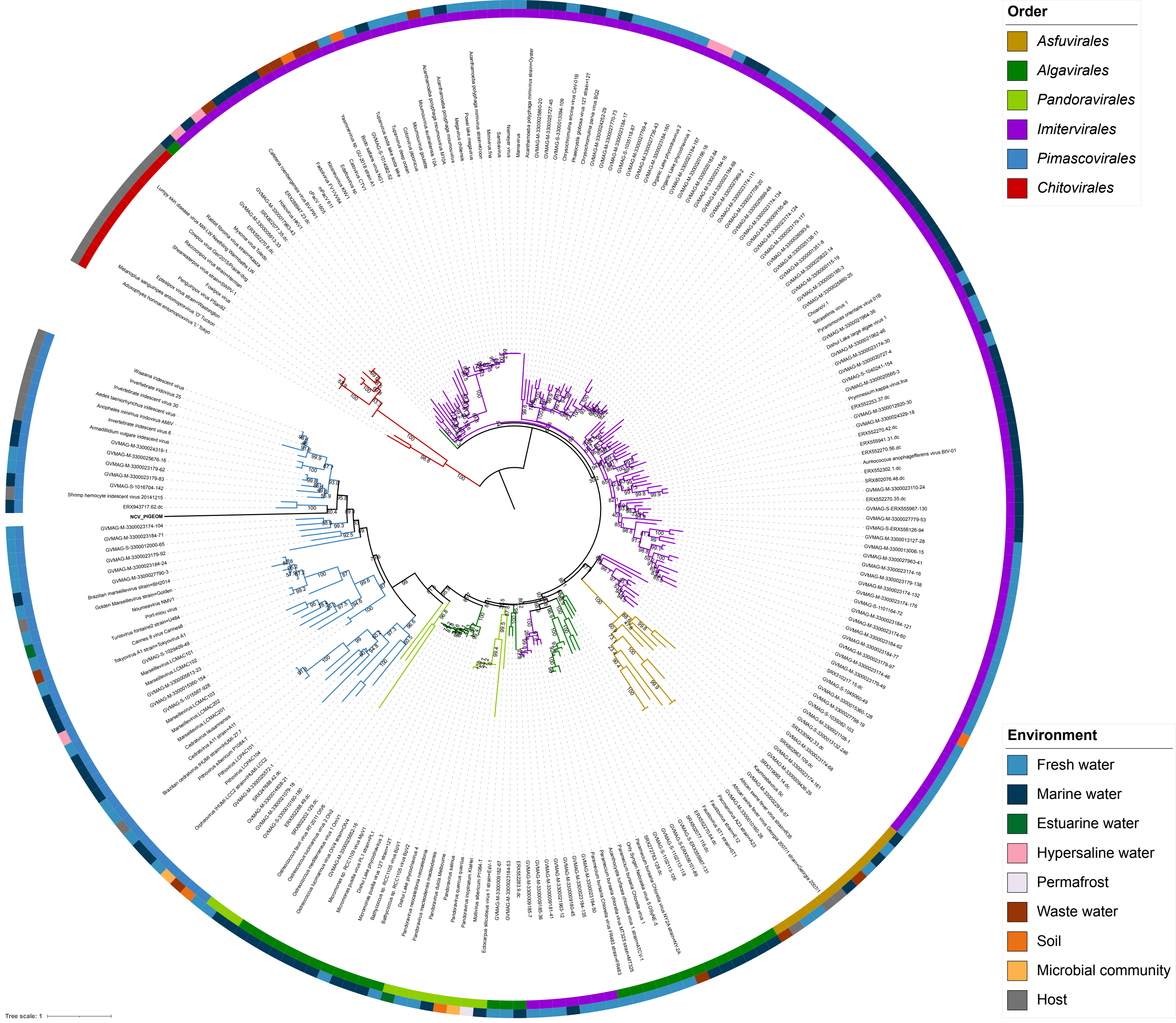


**Figure S2**. Maximum likelihood phylogenetic tree for *SFII* with a dataset containing 232 *Nucleocytoviricota*. The tree was constructed at the amino acid level. Branch values correspond to bootstrap support values. Branch colour coding and colours in the first ring around the tree correspond to the viral orders. *Chitovirales* was used as the outgroup. The colours in the second ring correspond to the environment in which the virus was found.

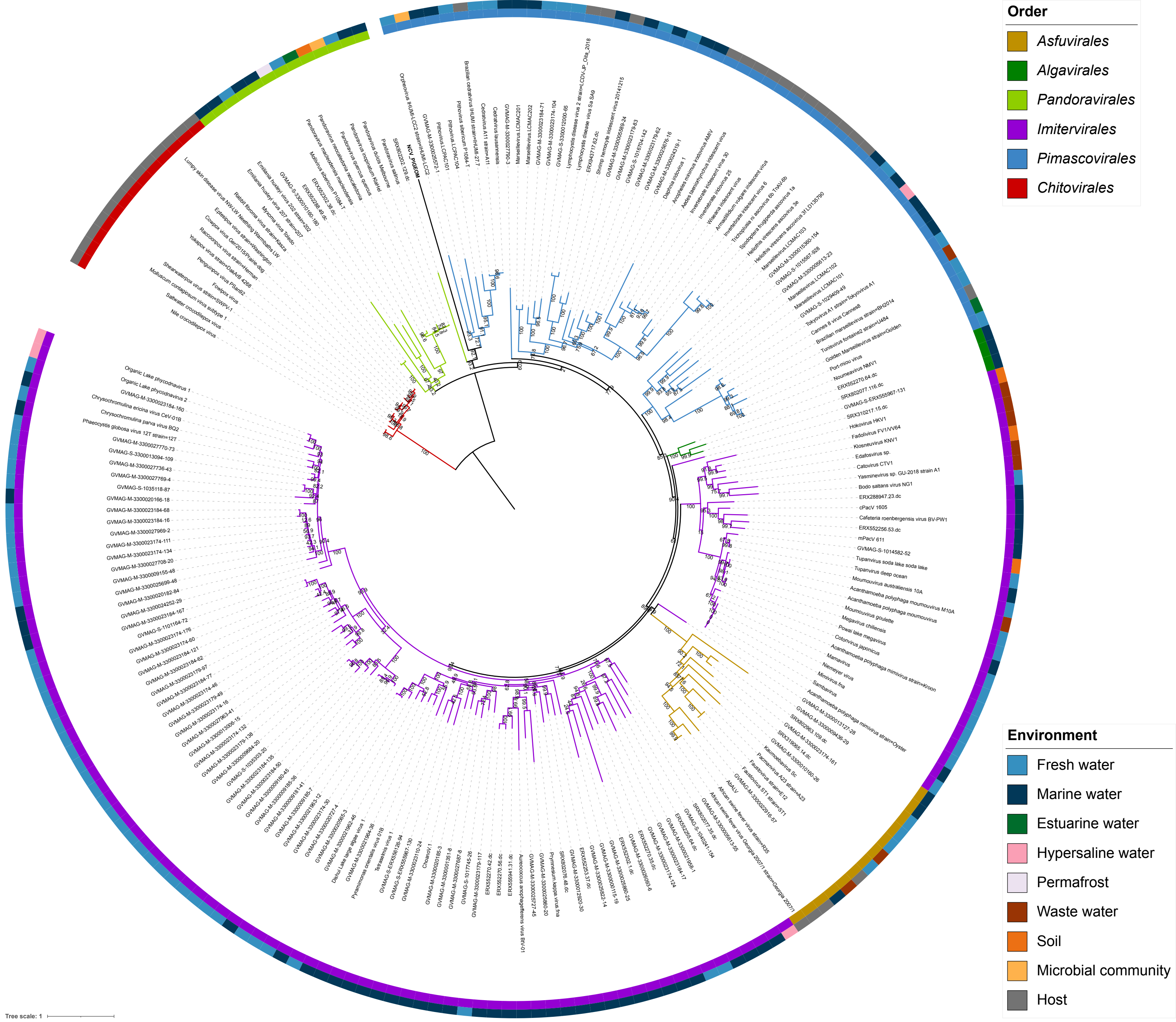


**Figure S3**. Maximum likelihood phylogenetic tree for *RNAPS* with a dataset containing 209 *Nucleocytoviricota*. The tree was constructed at the amino acid level. Branch values correspond to bootstrap support values. Branch colour coding and colours in the first ring around the tree correspond to the viral orders. *Chitovirales* was used as the outgroup. The colours in the second ring correspond to the environment in which the virus was found.

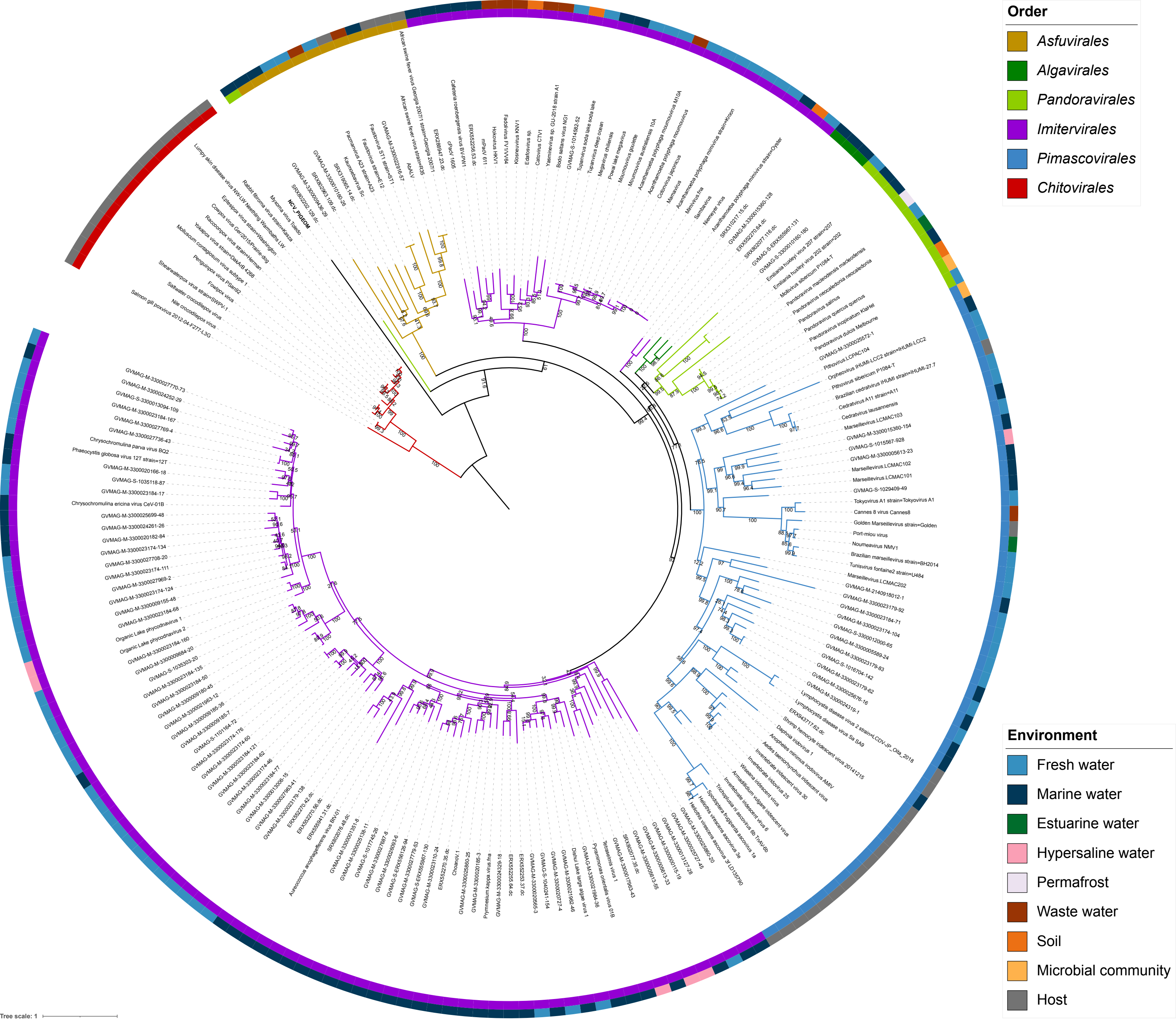


**Figure S4**. Maximum likelihood phylogenetic tree for *RNAPL* with a dataset containing 201 *Nucleocytoviricota*. The tree was constructed at the amino acid level. Branch values correspond to bootstrap support values. Branch colour coding and colours in the first ring around the tree correspond to the viral orders. *Chitovirales* was used as the outgroup. The colours in the second ring correspond to the environment in which the virus was found.

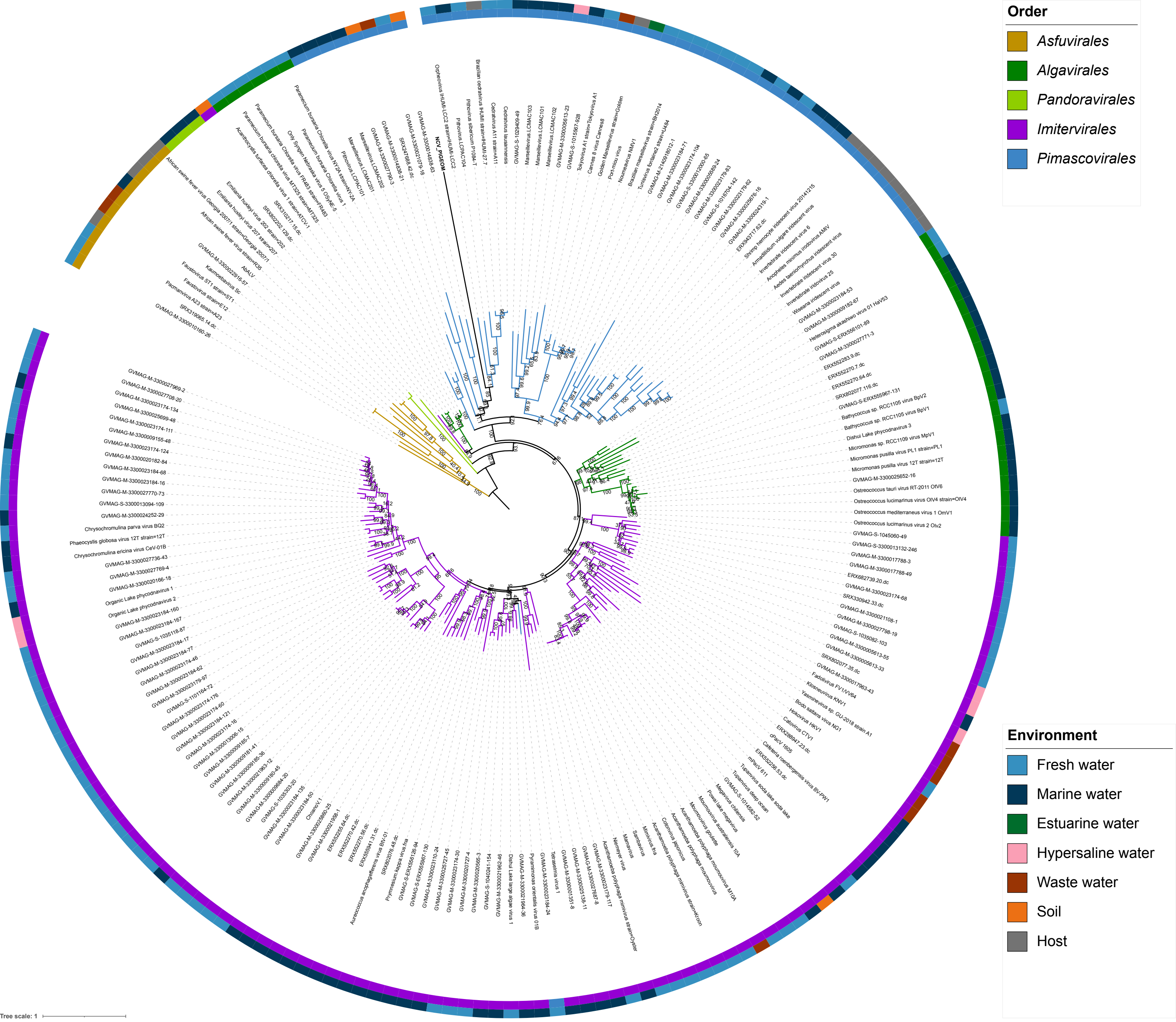


**Figure S5**. Maximum likelihood phylogenetic tree for *TopoII* with a dataset containing 201 *Nucleocytoviricota*. The tree was constructed at the amino acid level. Branch values correspond to bootstrap support values. Branch colour coding and colours in the first ring around the tree correspond to the viral orders. *Asfuvirales* was used as the outgroup. The colours in the second ring correspond to the environment in which the virus was found.

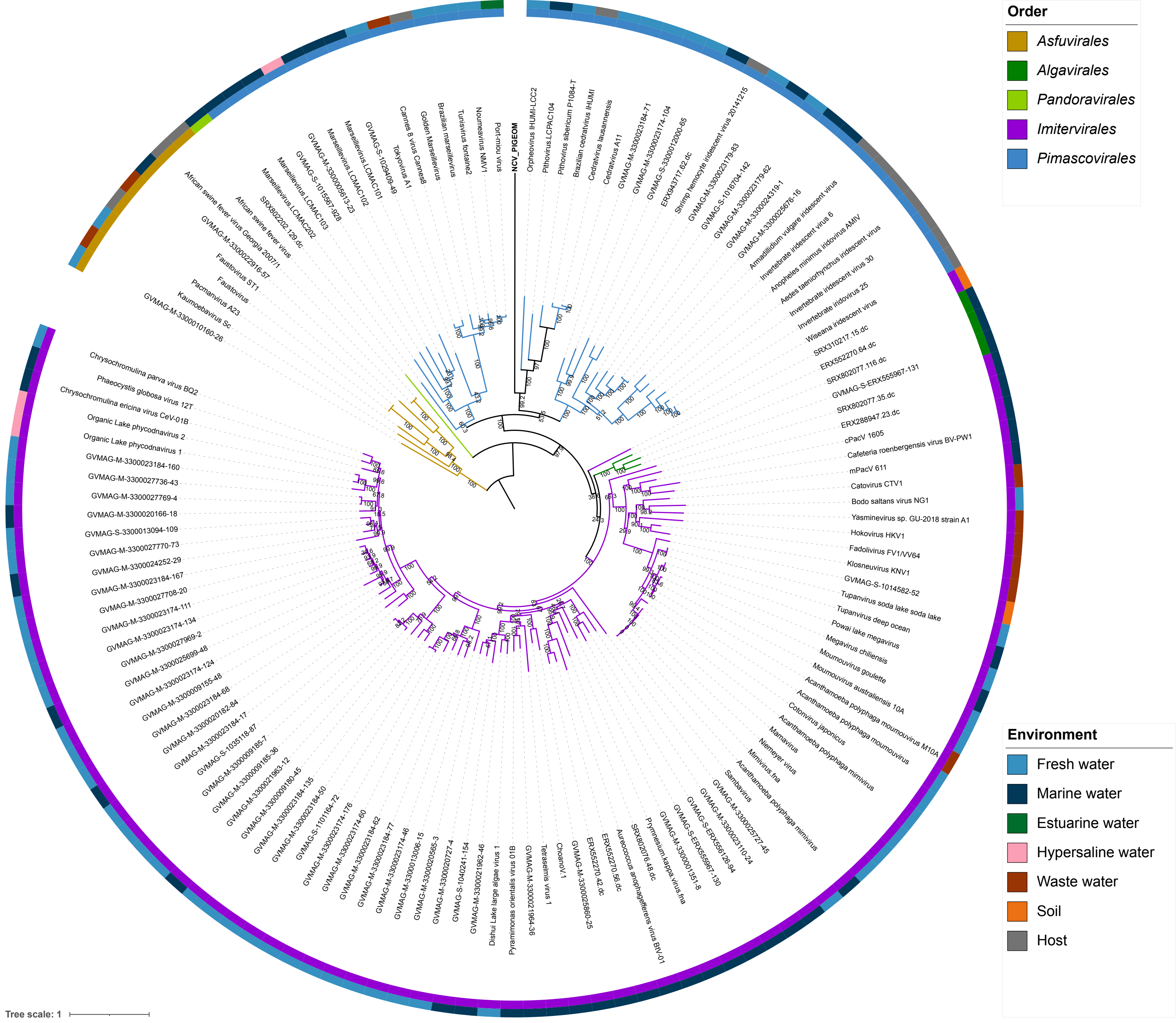


**Figure S6**. Maximum likelihood phylogenetic tree for a dataset containing 135 *Nucleocytoviricota*. The tree was constructed at the amino acid level from concatenated alignments of the *SFII*, *RNAPS*, *RNAPL*, *PolB*, and *TopoII* core genes. All 135 taxa contain all five core genes. Branch values correspond to bootstrap support values. Branch colour coding and colours in the first ring around the tree correspond to the viral orders. *Asfuvirales* was used as the outgroup. The colours in the second ring correspond to the environment in which the virus was found.
